# Clinicians’ management of patients potentially exposed to rabies in high-risk areas in Bhutan: A cross-sectional study

**DOI:** 10.1101/425884

**Authors:** Kinley Penjor, Nelly Marquetoux, Chendu Dorji, Kinley Penjor, Sithar Dorjee, Chencho Dorjee, Peter D Jolly, Roger S Morris, Joanna S. McKenzie

## Abstract

**Background:** Rabies is endemic in southern Bhutan, associated with 1–2 human deaths annually and accounting for about 6% of annual national expenditure on essential medicines. A WHO-adapted National Rabies Management Guidelines (NRMG) is available to aid clinicians in PEP prescription. An understanding of clinical practice in the evaluation of rabies risk in endemic areas could contribute to improve clinicians’ PEP decision-making.

**Methods:** A cross-sectional survey of clinicians was conducted in 13 health centers in high-rabies-risk areas of Bhutan during February–March 2016. Data were collected from 273 patients examined by 50 clinicians.

**Results:** The majority (69%) of exposure was through dog bites. Half the patients were children under 18 years of age. Consultations were conducted by health assistants or clinical officers (55%), or by medical doctors (45%), with a median age of clinicians of 31 years. Rabies vaccines were prescribed in 91% of exposure cases. The overall agreement between clinician’s rabies risk assessment and the NRMG for the corresponding exposure was low (kappa =0.203, p<0.001). Clinicians were more likely to underestimate the risk of exposure than overestimate it. Male health assistants were the most likely to make an accurate risk assessment and female health assistants were the least likely. Clinicians from district or regional hospitals were more likely to conduct accurate risk assessments compared to clinicians in Basic Health Units (Odds Ratios of 7.8 and 17.6, respectively).

**Conclusions:** This study highlighted significant discrepancies between clinical practice and guideline recommendations for rabies risk evaluation. Regular training about rabies risk assessment and PEP prescription should target all categories of clinicians. An update of the NRMG with more specific criterions for the prescription of RIG might contribute to increase the compliance, along with a regular review of decision-making criteria to monitor adherence to the NRMG.

**Author summary:** Human rabies remains an important public health threat in Bhutan, especially in southern regions where canine rabies is endemic. The steady increase in number of patients reporting to hospitals following dog bites means escalating costs of post-exposure prophylaxis for the country. We investigated attitudes and practices of clinicians who manage patients with potential rabies exposure, in the endemic area. The risk of rabies exposure in the study area is mostly associated with dog bites, involving children half the time. Rabies vaccines were prescribed in 9 out of 10 exposure cases, while immuno-globulins were rarely prescribed. The study confirmed the perceived lack of compliance of clinicians with guideline recommendations for assessing rabies risk. This results in under-estimating the rabies risk in potentially risky exposures in high-rabies-risk areas. Our work underscore the importance of targeted training of female health assistants, doctors, and clinicians in basic health units to improve the management of rabies exposure. In particular there is need to update the national guidelines regarding indications and use of rabies immune-globulins.

## Introduction

In Asia, rabies remains a major public health threat, with an estimated 39,000 deaths annually, mostly due to spill over from the canine reservoir [1]. Wider use of post-exposure prophylaxis (PEP) might reduce human mortalities in this region of the world [2]. On the other hand, the escalating cost of life-saving PEP represents a major burden to both national economies and families, mostly in poor rural communities [1, 3, 4].

In Bhutan, the number of reported animal rabies cases was stable in the decade 1996 to 2005 but increased during 2006 to 2008, mostly in 4 districts in southern Bhutan bordering India [5]. Maintenance of rabies in the canine reservoir in the south Bhutan was likely due to low coverage of dog vaccination programs. Since 2009, mass dog sterilization and vaccination programs contributed to a decline in the incidence of canine rabies [6]. However, the disease remains endemic in South Bhutan and in some pockets in East Bhutan. All 13 human deaths due to rabies recorded between 2009 and 2017 came from southern districts where the estimated average annual incidence is 0.4 deaths/100,000 population [7, 8]. Rabies PEP (wound treatment, vaccination with or without immunoglobulin administration) is provided free of charge for the public by the government of Bhutan. Recent estimates show that current PEP intervention effectively averts about 15 human deaths annually in rabies endemic areas [9]. However, PEP prescription without a thorough rabies exposure risk assessment results in substantial costs to the health sector [10]. Between 2009 and 2016, the annual number of dog bite patients increased from 1000 to over 7000 [7]. As per the Ministry of Health record, an estimated annual cost of PEP in Bhutan is about Nu. 8.5 million (USD 131,000) – approximately 6% of the essential medicines budget (S1)

To assess the rabies risk in potentially exposed patients, the Ministry of Health in Bhutan recommends clinicians follow the National Rabies Management Guidelines (NRMG) [11]. The guidelines are in line with WHO recommendations. The rabies risk assessment in endemic areas should be based on the nature of exposure to animals or their products and the nature of the injury. Prescription of anti-rabies vaccine (ARV) is recommended for patients with a moderate or severe risk (NRMG Categories 2 and 3), while ARV associated with rabies immunoglobulin (RIG) are recommended for Category 3 exposures. However, a shortage of RIG supply in Bhutan resulted in RIG treatment being reserved for patients with exposure to suspected or confirmed rabid animals only. The current wording in the NRMG around RIG prescription lacks clarity and does not provide clear direction for clinicians.

Despite the availability of the NRMG, there is public concern that both sporadic human deaths due to rabies and rising PEP expenditure could be results of inappropriate rabies PEP prescription by clinicians. However, there is currently no published evidence supporting this assumption. Therefore, this study investigated rabies risk assessment and PEP prescription practices in potentially exposed patients by clinicians in the high-rabies-risk areas of southern Bhutan. The aim was to provide a better understanding of clinician’s knowledge and practices with respect to managing patients potentially exposed to rabies and identify measures to improve these where necessary to strengthen the national effort to reduce the burden of rabies.

## Materials and Methods

We conducted a cross-sectional study to assess clinicians’ rabies risk assessment process and PEP prescription decisions during the management of patients potentially exposed to rabies. The study was approved by the Research Ethics Board of Health, vide approval ref.no. *REBH/Approval/2016/001*.

### Study sites and participants

All 13 health centers with doctors in the medical staff, i.e. hospitals and grade I Basic Health Units (BHU-I), located in the high rabies-risk belt of southern Bhutan were included in the study (Fig 1). All clinicians involved in the management of patients potentially exposed to rabies infection from animals were included. The term ‘clinician’ in this study refers to staff who treat patients, including doctors and paramedical staff (clinical officers and health assistants).

**Fig 1.**
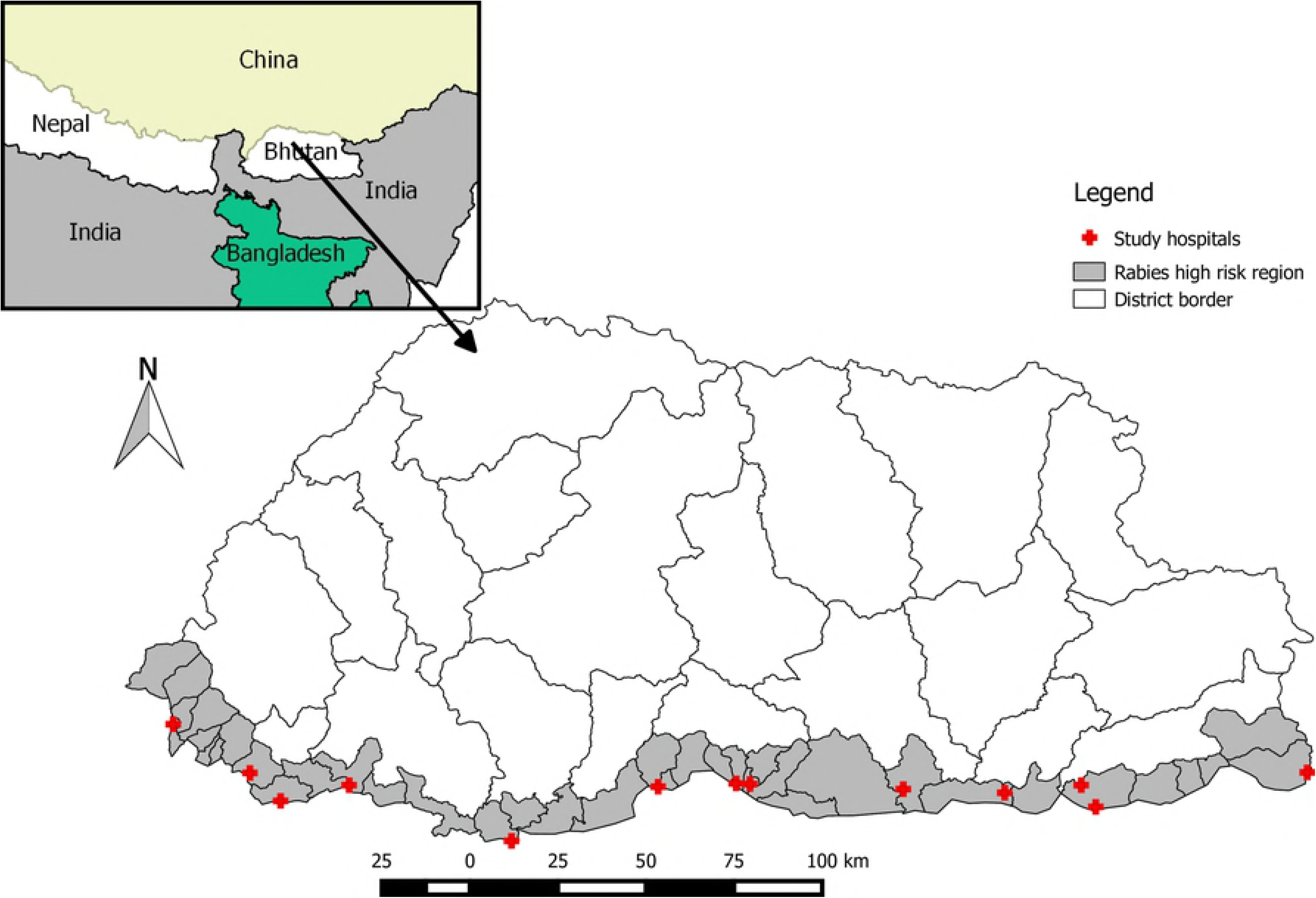
Map of Bhutan showing rabies high-risk region and the location of the 13 study sites (Map created using open source QGIS 2.12 software downloaded from http://monde-geospatial.com/download-qgis-2-12-lyon-the-last-released-version-of-qgis/ and shapes files from https://gadm.org/download_country_v3.html)

### Study design and data collection

The study was conducted from 1^st^ February to 31^st^ March 2016. Cases were defined as all patients presenting to a hospital or BHU-I for treatment following potential exposure to rabies from an animal or animal product. Cases were identified prospectively and a questionnaire was completed by designated, specifically trained staff in each study site, while observing the patient’s consultation with the clinician. Additional information or verification was sourced directly from the clinician or from the patient where adequate information was not collected by the clinician during the consultation. For example, the patient was interviewed about the exposure event after the consultation if the clinician did not elicit the required information. The data collected included demographic information about the patient, clinicians’ level of qualification and years of experience, whether the clinician used a pre-determined set of epidemiological questions about the nature of the animal exposure, the patient’s response, the rabies risk as assessed by the clinician, and whether PEP was prescribed. Information was cross-validated from copies of the PEP case record sheets maintained by hospital staff.

### Data analysis

#### Demographics of clinicians and patients

Summary statistics were calculated to describe the population of clinicians performing clinical assessments of patients potentially exposed to rabies, and the demographics of the patient population included in the study. Given individual clinicians managed a varying number of cases potentially exposed to rabies, summary statistics for clinician demographics were weighted according to the number of such cases managed by each clinician. All personal identifiable information of both patient and clinician were removed to anonymize and protect their privacy before the analysis of data was conducted

#### Types of exposure

The different types of potential exposure and the animal species involved were determined from each patient’s account of the exposure event. Exact binomial tests were used to test for equi-probability of males versus females for various types of exposure.

#### Rabies risk assessment conducted by clinicians

Our questionnaire comprised a set of 23 epidemiological questions to evaluate the level of rabies risk in a patient exposed to a potentially infected animal. The questionnaire was prepared using the NRMG and a rabies expert panel and covered date, type of exposure, animal species involved, vaccination status (dog and cat), potential rabies status of the animal and past PEP history of the patient.

We identified sets of relevant epidemiological questions for three types of exposures, namely: direct exposure to an owned animal or a stray animal, and indirect exposure to any animal through contact with animal products or fomites. For example, questions about indirect exposure were irrelevant for direct exposure cases. Similarly, questions about vaccination status of the animal were irrelevant for patients bitten by a stray animal.

Firstly, we described the proportion of clinicians asking the relevant epidemiological questions pertaining to cases with each type of exposure. Secondly, we independently classified each case into one of three rabies risk categories (none, moderate, severe) by comparing the epidemiological information provided by the patient, either during or following the consultation, with the current NRMG [11]. The criteria that we used for rabies risk classification are presented in Table 1.

**Table 1.** Criteria for rabies risk assessment and recommended PEP prescription in case of exposure to “suspect or rabid animals”, extracted from the WHO adapted National Rabies Management Guidelines (2014) in Bhutan

| Exposure type | Risk category | Recommended PEP |
| --- | --- | --- |
| Licks on intact skin, touching, feeding of animals.<br><sup>1</sup>Consumption of butter, curd, cheese, whey (dachu), cooked meat.<br>Petting, bathing or coming in contact with utensils used on a suspected rabid animal. | None (Category 1) | Not recommended, if reliable case history available. |
| Person consuming unboiled or unpasteurized milk, buttermilk, | Moderate (Category 2) | Wound management |
| uncooked meat from rabid animal.<br>Nibbling of uncovered skin by suspected rabid animal.<br>Minor scratches or abrasions without bleeding.<br>Person who handles or prepares meat or handles carcass of rabid animals. |  | Provide anti-rabies vaccine immediately<br>Stop vaccination if animal remains healthy throughout the observation period of 10 days or if the animal is proven to be negative for rabies by a reliable laboratory using an appropriate diagnostic technique |
| Single or multiple transdermal bites or scratches<br>Licks on broken skin<br>Contamination with mucous membrane with saliva (i.e. licks or splash on oral cavity, eyes, nose, external genitalia) | Severe (Category 3) | Wound management<br>Provide anti-rabies vaccine immediately<br>Provide rabies immunoglobulin<br>Animal observation and management as above (Category 2) |

For patients for whom the clinician assessed the rabies risk, we estimated the agreement between the risk category assigned by clinicians and the independently assessed risk based on the NRMG, using the weighted Cohen’s kappa statistic for ordinal variables and equal weights. Thirdly, we performed logistic regression to evaluate potential factors associated with “agreement” between the clinician and the NRMG risk assessment. A binary variable “agreement” (0: disagreement between clinician and the NRMG, 1: agreement between clinician and the NRMG) was created. Variables evaluated in the bivariate model were:

- gender;
- professional experience (number of years);
- designation of the clinician: medical doctor, assistant clinical officer, health assistant;
- highest level of qualification: Bachelor of Medicine, Bachelor of Surgery (MBBS), Diploma or Certificate;
- health center type: Basic Health Unit, district hospital, regional referral hospital;
- district (9 levels);
- actual level of risk of the exposure event according to the NRMG (none, moderate, severe).

Variables significant at P<0.3 were used to fit a multivariate model. Observations were clustered by clinician, hospital and district, hence these 3 variables were used as nested random effects in the model. We used a stepwise backward model selection process using the lowest Akaike Information Criterion (AIC). Potential interactions between fixed effect variables in the final model were evaluated and selected using the lowest AIC.

### Assessment of PEP prescription by the clinician

We described the clinicians’ practices in relation to the prescription of PEP, in particular ARV and RIG, according to rabies risk classification as assessed by clinicians and independently assessed according to the NRMG. All analyses were performed using R [12].

## Results

Fifty clinicians participated in this study and questionnaires were completed for 273 patients. Each patient had only one consultation for potential rabies exposure during the study period and each clinician saw an average of 5.5 such patients (median: four patients, range 1-19). Most consultations occurred in district hospitals (177/273, 64.8%), 59 in BHU-I and 37 in regional hospitals.

### Clinicians’ demographics

All clinician level statistics were weighted by the number of consultations provided by each clinician. The median age of clinicians was 31 years (range 25 to 55 years) and median duration of clinical experience was 6 years (range 1 to 35 years). Doctors had a median of two years clinical experience while clinical officers and health assistants had a median of 22 and 17 years, respectively. Health assistants or clinical officers who held a certificate or diploma qualification conducted the majority of consultations (55%) while MBBS doctors conducted the rest. Male staff performed nearly three quarters of the consultations (Table 2).

**Table 2.** Demographic features of clinicians who led consultations for 273 patients potentially exposed to rabies in high risk areas in Bhutan during the study period.

| Variable | Freq. | Percent. |
| --- | --- | --- |
| **Designation** |  |  |
| Medical Doctor | 124 | 45.4 |
| Clinical Officer (or Assistant CO) | 34 | 12.5 |
| Health Assistant | 115 | 42.1 |
| **Gender** |  |  |
| Males | 201 | 73.6 |
| Female | 72 | 26.4 |
| **Type of Health Center** |  |  |
| District Hospitals | 177 | 64.8 |
| Regional Hospitals | 37 | 13.6 |
| Basic Health Units | 59 | 21.6 |

### Patient demographics

Patients’ age spanned 2 to 85 years, with half under 18 years of age (137/273). The frequency of potential rabies exposure in children decreased regularly with age after 2 years of age; children <2 years of age were infrequently exposed (**Fig 2**). Of the patients presenting, 97% were Bhutanese nationals; 55% were male, and 45% were female (P>0.05). Half of the patients were preschoolers or students, and 16% were farmers (Fig 3). The majority of patients (208/267, 78%) presented to the health center or hospital on the day of exposure or the following day.

**Fig 2.**
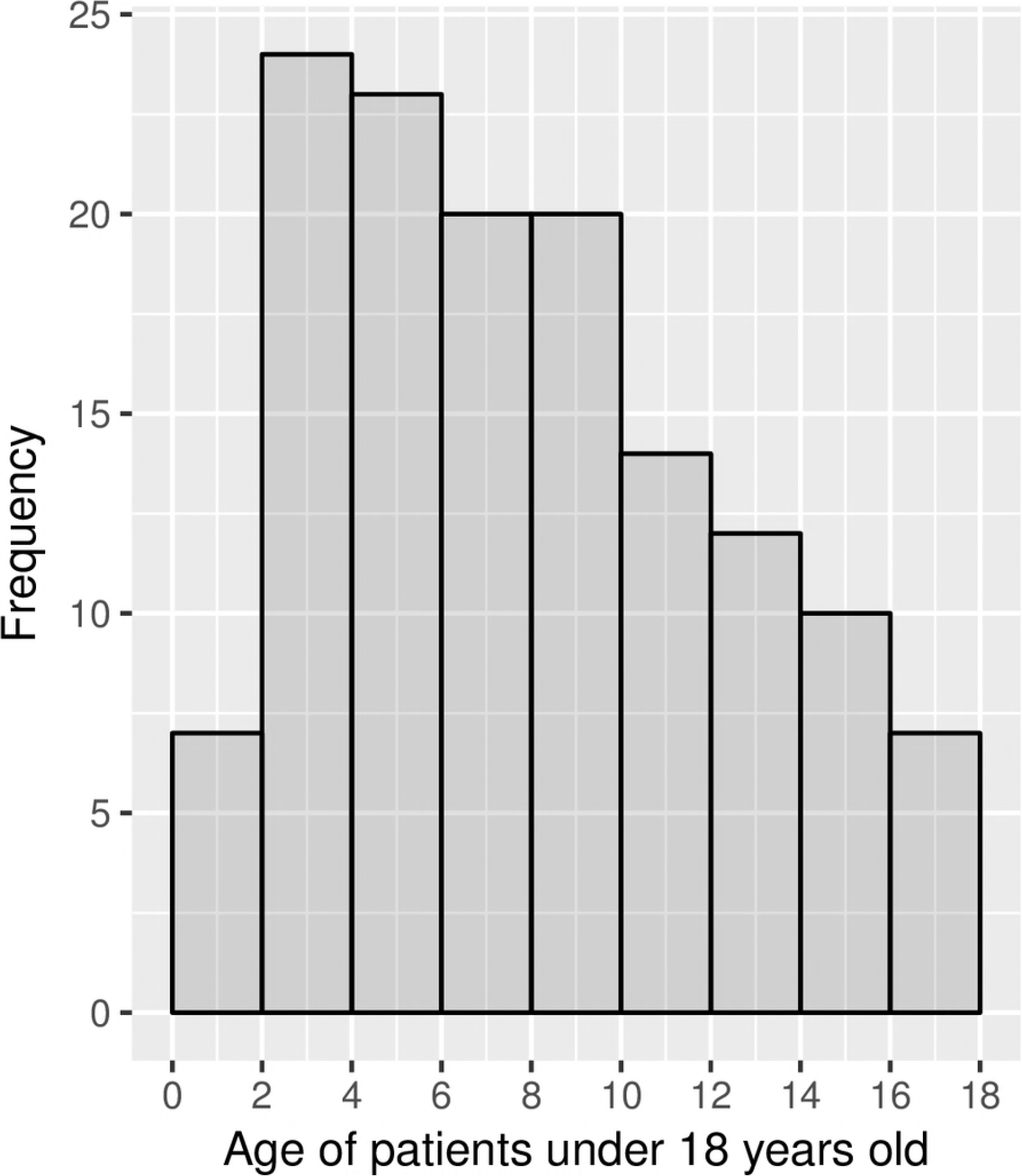
Age distribution of patients under 18 years old (n=137) seeking treatment for potential exposure to rabies in the study centers during the period 1st February to 31st March 2016

**Fig 3.**
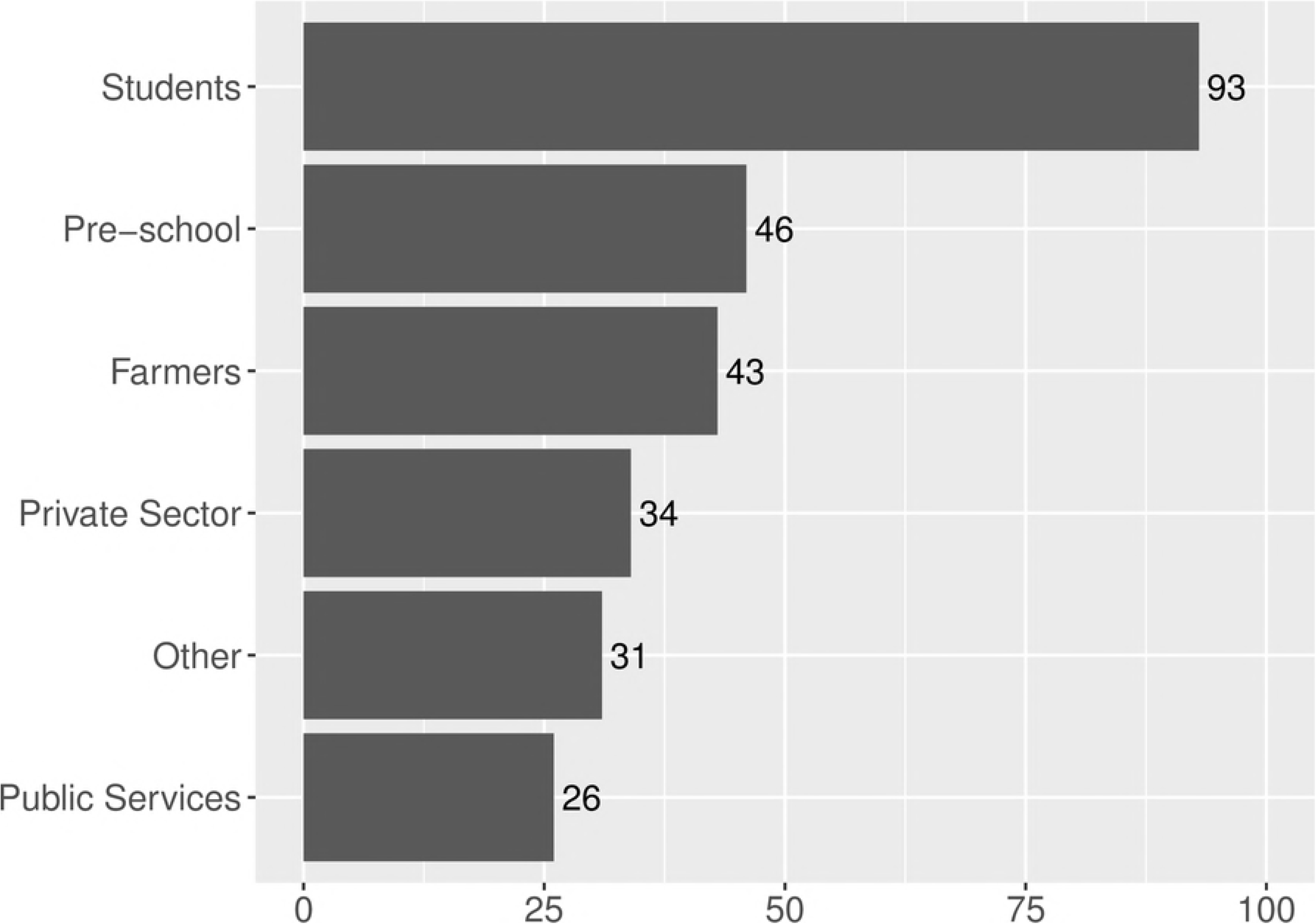
Occupation of 273 patients presenting for treatment following potential exposure to rabies in high-risk areas in southern Bhutan

### Types of exposure

The majority of patients were potentially exposed to rabies as a result of dog bites (189/273, 69%). Among dog bites, 67 (35%) were inflicted by free-roaming (also referred to as ‘stray’) dogs and 123 (65%) by pet dogs. There was no significant difference between the proportions of male or female patients for each of the exposure categories presented in Table 3.

**Table 3.** Animal species involved, type of exposure and demographics for 273 patients potentially exposed to rabies through contact with animals

| Variable | Frequency (%) | Patient demographics |  |  |
| --- | --- | --- | --- | --- |
|  |  | Male | Female | Mean age |
| Animal species |  |  |  |  |
| Pet dog | 140 (51.3 %) | 79 | 61 | 23.3 |
| Free roaming dog | 81 (29.7 %) | 48 | 33 | 20.5 |
| Cat | 40 (14.7 %) | 16 | 24 | 22.4 |
| Cattle/buffalo | 12 (4.4 %) | 5 | 7 | 34.8 |
| Rodents/wild animals | 5 (1.8 %) | 2 | 3 | 26 |
| Exposure types |  |  |  |  |
| Bites (with bleeding) | 152 (55.7 %) | 85 | 67 | 23.7 |
| Bites (no bleeding) | 61 (22.3 %) | 35 | 26 | 22.9 |
| Scratches | 52 (19.1 %) | 26 | 26 | 17.3 |
| Licks or Nibbles | 11 (4 %) | 5 | 6 | 18.3 |
| Carcass handling (cattle/buffalo) | 6 (2.2 %) | 3 | 3 | 38.2 |
| Indirect exposure (consumption of milk/milk products or contact with animal products) | 8 (2.9 %) | 3 | 5 | 26.0 |

### Rabies risk assessments

In nearly all consultations, the clinicians investigated the type of rabies exposure by asking relevant questions during the consultation. The exact type of exposure, the date, the wound site and the species involved were obtained in over 95% of clinical assessments. Details of the epidemiological information that clinicians collected to assess rabies risk are detailed in (Fig 4). The risk profile of the 272 patients for whom information was available to independently classify rabies risk according to the NRMG was 57% severe risk, 43% moderate risk and only 1 (0.3%) was no risk.

**Fig 4.**
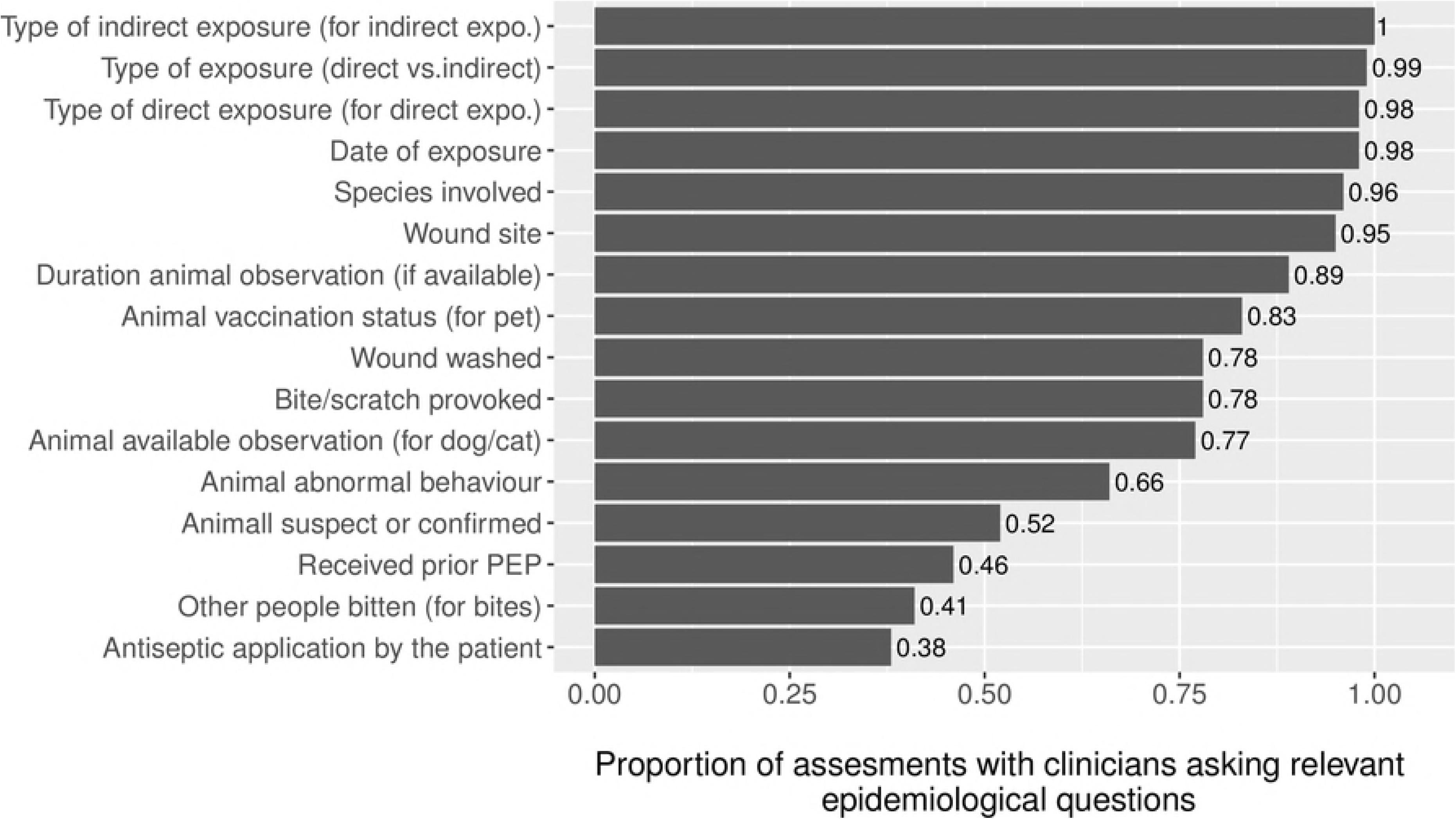
Proportion of relevant epidemiological questions asked by the clinicians for each exposure type (indicated in brackets); the denominator varied between 8 and 273, depending on the frequency of the type of exposure

The attending clinicians recorded the corresponding rabies risk assessment (none, moderate, severe) on the patient’s sheet for 194 (71%) of the 273 patients. Of these 194 risk assessments performed by clinicians, only 102 (53%) were correctly classified when compared to the NRMG (Table 4). It was more frequent for clinicians to underestimate rabies risk (37%) than to overestimate it (11%). Of the 154 cases independently classified as severe rabies risk, 62 (40%) were correctly classified by clinicians, 45 (29%) were classified as moderate risk, 9 (6%) were classified as having no rabies risk and 38 (25%) were not classified or recorded by the clinicians. Based on results in Table 4, the kappa agreement test statistics was 0.203 (p<0.001), indicating a poor agreement of rabies risk categorization between the clinicians and the NRMG.

**Table 4.** Comparison of clinicians’ classification of rabies risk versus National Rabies Management Guidelines (NRMG) for 272^1^ patients potentially exposed to rabies

| Risk category (NRMG): | Risk category assigned by the clinician: |  |  |  |
| --- | --- | --- | --- | --- |
|  | None | Moderate | Severe | NA<sup>2</sup> |
| None | **0** | 1 | 0 | 0 |
| Moderate | 17 | **40** | 20 | 40 |
| Severe | 9 | 45 | **62** | 38 |

<sup>1</sup> One exposure event could not be categorized, even retrospectively, due to missing data
<sup>2</sup> Not Available: risk either not assessed or not recorded by the clinician

Potential explanatory variables associated with the agreement between the clinician’s risk assessment and the NRMG (final model) are presented in Table 5. The effect of clinical designation interacted with that of gender. Male health assistants were the most likely to make an accurate risk assessment and female health assistants were the least likely. Female and male doctors were not significantly different. Male health assistants were three times more likely to make an accurate risk assessment than male doctors, whereas female doctors were twice as likely to be accurate than female health assistants. Male health assistants were 12 times more likely to make an accurate risk assessment than female health assistants. Clinicians from district or regional hospitals were much more likely to conduct accurate risk assessments compared to clinicians in Basic Health Units (Odds Ratio of 7.8 and 17.6, respectively), independent of the clinician’s designation in the different healthcare facilities. The random effects (clinicians nested in hospitals nested in districts) suggested that after taking into account the variation in assessment accuracy associated with clinicians, hospitals and the fixed effects, there was no residual variation between districts.

**Table 5.**
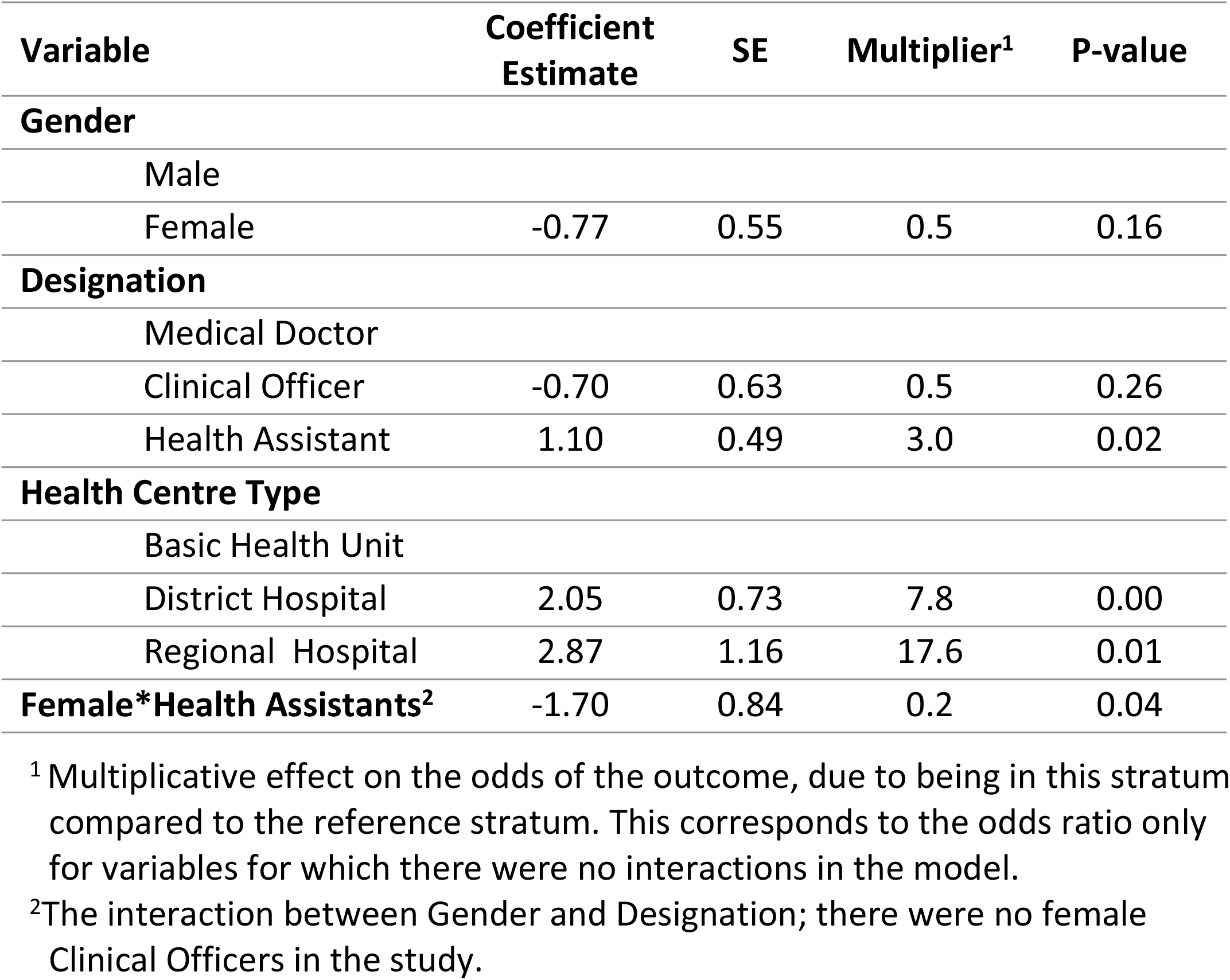
Final multivariate logistic regression model of factors associated with making an accurate rabies exposure risk assessment, defined by agreement with the NRMG.

| Variable | Coefficient Estimate | SE | Multiplier<sup>1</sup> | P-value |
| --- | --- | --- | --- | --- |
| **Gender** |  |  |  |  |
| Male |  |  |  |  |
| Female | -0.77 | 0.55 | 0.5 | 0.16 |
| **Designation** |  |  |  |  |
| Medical Doctor |  |  |  |  |
| Clinical Officer | -0.70 | 0.63 | 0.5 | 0.26 |
| Health Assistant | 1.10 | 0.49 | 3.0 | 0.02 |
| **Health Centre Type** |  |  |  |  |
| Basic Health Unit |  |  |  |  |
| District Hospital | 2.05 | 0.73 | 7.8 | 0.00 |
| Regional Hospital | 2.87 | 1.16 | 17.6 | 0.01 |
| **Female\*Health Assistants<sup>2</sup>** | -1.70 | 0.84 | 0.2 | 0.04 |

<sup>1</sup> Multiplicative effect on the odds of the outcome, due to being in this stratum compared to the reference stratum. This corresponds to the odds ratio only for variables for which there were no interactions in the model.
<sup>2</sup> The interaction between Gender and Designation; there were no female Clinical Officers in the study.

### Clinicians’ PEP prescription practices

The number of patients for whom clinicians prescribed ARV by rabies risk category as assessed by the clinician and as independently assessed according to the NRMG is shown in Table 6. Neither ARV nor RIG was prescribed for 1 patient assessed by the clinician to be severe risk and ARV was not prescribed for a second patient assessed to be moderate risk. Conversely, clinicians prescribed ARV for 10 (38%) of 26 patients whom they assessed as having no rabies risk and 75 (95%) of 79 patients for whom they did not assess rabies risk. Considering the independent risk categorization of patients according to NRGM, 7 (5%) of 154 severe risk and 16 (14%) of 117 moderate risk patients were not prescribed ARV. PEP and other treatments prescribed by the clinicians in this study are described in (Fig 5).

**Table 6.** Number of patients prescribed ARV by risk category of potential rabies exposure as assessed by clinicians and as independently assessed according to the NRMG.

| Rabies risk: | According to the clinician: |  |  |  | According to NRMG: |  |  |
| --- | --- | --- | --- | --- | --- | --- | --- |
|  | None | Mod. | Severe | NA | None | Mod. | Severe |
| ARV prescription: |  |  |  |  |  |  |  |
| No ARV | 16 | 2 | 1 | 4 | 0 | 16 | 7 |
| ARV | 10 | 84 | 81 | 75 | 1 | 101 | 147 |
| Additional RIG* | 0 | 1 | 2 | 0 | 0 | 1 | 2 |

\*3 patients were prescribed RIG on top of ARV: one patient handling the carcass of a confirmed rabies cow, misclassified as risk 3 category, one patient bitten by a suspect rabies dog, misclassified as risk 2 but prescribed RIG, one patient bitten by a suspect rabies dog, correctly classified as risk 3.

**Fig 5.**
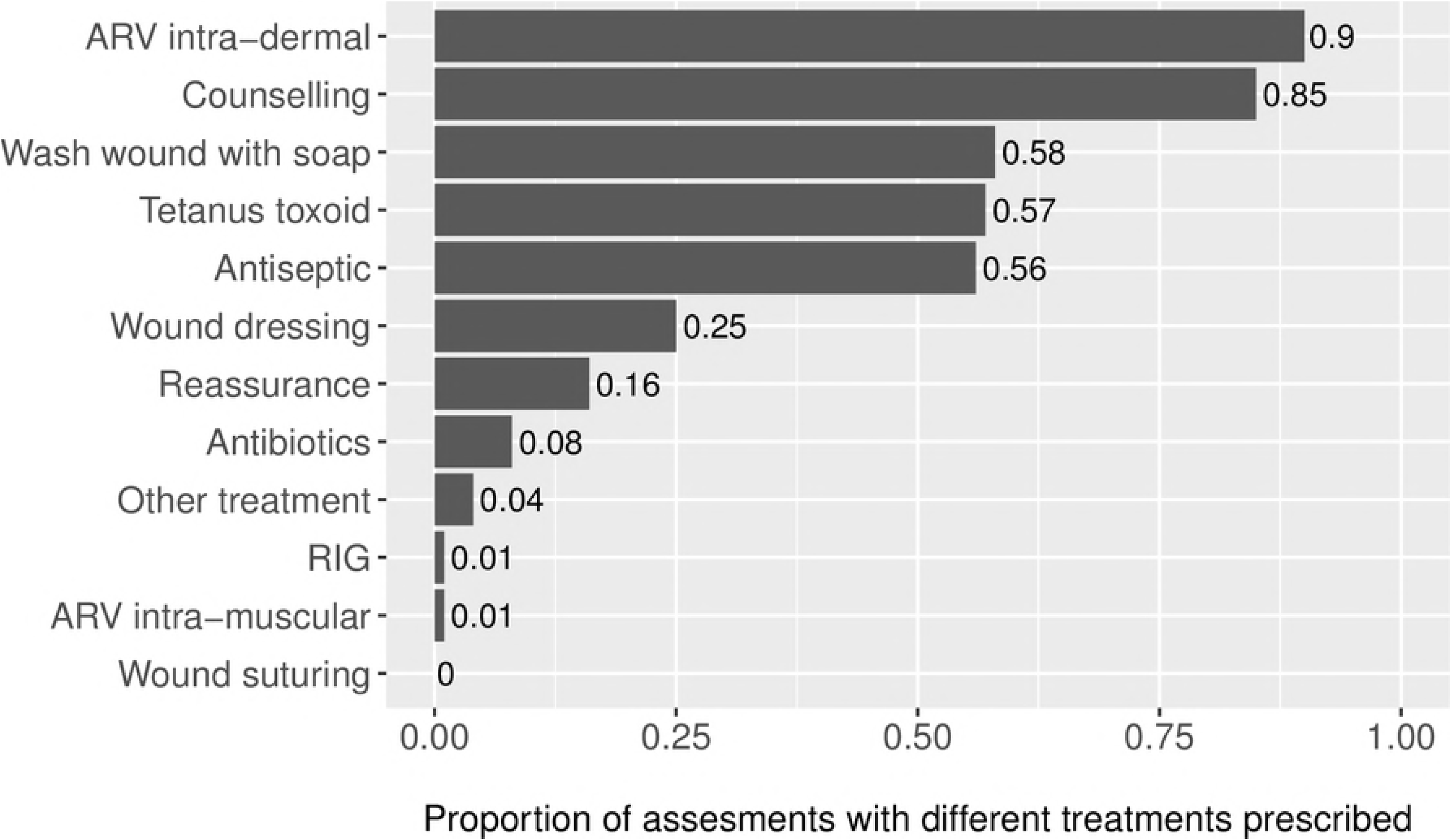
Proportion of different therapeutic measures prescribed by the clinicians; the denominator varies between 257 and 273 after removing some missing information

## Discussion

This study was the first attempt to describe and evaluate clinical practices in the management of patients potentially exposed to rabies through contact with animals, in high-rabies-risk areas of Bhutan. We described and analyzed consultations for 273 patients who were potentially exposed to rabies, which were conducted by 50 clinicians in 13 health centers in the study area. Most consultations were conducted by health assistants and clinical officers (55%), and the remainder by medical doctors (45%), mostly in district hospitals (65%). Most consultations were provided by junior health workers, with a median clinical experience of six years.

Dog bites constituted the main source of potential rabies exposure (69%) and half of the patients were under 18 years old. The frequency of potential rabies exposure in children decreased with age, except for children under two years old, which were rarely presented. However, the age distribution of the underlying study population was not taken into account, neither was the possible differential bias of reporting to health centers following potential rabies exposure in adults versus children. Age-specific risks thus cannot be inferred from these data.

We attempted to minimize information biases from the clinicians through prior communication. Observational biases from the interviewer were mitigated by training on information recording. However, clinicians could have been influenced towards more rigorous risk assessment during the study, so the results likely over-estimate the performance of clinicians.

### Risk classification

The independent rabies risk classification using the NRMG guidelines for 272 of the 273 cases for whom information was available showed 57% of cases had a severe risk, 43% a moderate risk and only 1 case was classified as having no risk. Clinicians collected epidemiological information about the type of exposure and species involved for almost all cases. However, a rabies risk assessment was performed and recorded for only 194 (71%) of cases. Clinicians’ risk categorization for this group showed a low level of agreement with the NRMG (kappa = 0.203), and a tendency for clinicians to underestimate exposure risk.

Male health assistants were most likely to make an accurate risk assessment and female health assistants were the least likely. Female and male doctors were not significantly different. Male Health Assistants were more likely to accurately assess risk than male doctors (Odds Ratio=3), whereas female doctors were more likely to be accurate than female Health Assistants (Odds Ratio =1.9). In addition, male Health Assistants were more likely to assess rabies exposure risk according to the guidelines than female Health Assistants (Odds Ratio = 12). Clinicians from district or regional hospitals were more likely to conduct risk assessments in agreement with the NRMG than clinicians in Basic Health Units (Odds Ratios of 7.8 and 17.6, respectively).

These results possibly reflect the availability of opportunities and participation by clinicians in capacity building training programs for rabies and PEP conducted by the Ministry of Health as was observed in a Haitian study [13]. The attitude of clinicians towards training might differ depending on their level of qualification. Health assistants are more readily available to attend such locally organized trainings. Most doctors, however, are unable to attend or oblivious to district training sessions. The apparent better performances of male health assistants compared to male doctors may reflect the successful impact of such continuing education opportunities, rather than initial education level. The lower level of agreement between risk assessments conducted by female Health Assistants and the NRMG than those conducted by male Health Assistants may reflect lower access or uptake of professional training by women compared to their male counterparts. In addition, male health assistants were more experienced (median of 29 years of clinical experience) compared to male or female doctors (median of 2 years of experience) and female health assistants (median of 10 years of experience). Due to this correlation between clinical designation and the number of years of experience, both variables could not be in the final model and the best fitting model was the model with clinical designation and without clinical experience.

Better performance of clinicians in hospitals compared to basic health units might be associated with lack of training and awareness of the guidelines by BHU level staff. This might also partially reflect the effect of clinical experience, even though the coefficients for type of health center was virtually unchanged when adding this variable to the model, after accounting for designation. Nevertheless, junior clinicians are often posted in lower level health facilities rather than hospitals, as per government policies. Moreover, the majority of rabies assessments (78%) occurred in hospitals rather than basic health units, hence junior clinicians in basic health units may be comparatively less experienced.

Specific rabies risk assessment and PEP training should target all clinicians involved in managing cases potentially exposed to rabies, including doctors, since the latter is equally in the frontline and tend to not perform as well as trained Health Assistants (particularly males), similar to Indian study conducted in eight cities [14]. Gender parity in the training of health professionals should also be pursued to ensure the engagement of female clinicians, as well as targeted training of staff in basic health units, as this is likely to make an important difference in improving the accuracy of rabies assessments.

According to the NRMG, a rabies risk assessment is recommended in cases exposed to “suspected or confirmed rabid animals”. However, there is no clear definition for a “suspected” rabies case in animals. In fact, since rabies is considered endemic in southern Bhutan, all potential vector animals involved in an exposure event should be considered as suspected rabies and followed by a risk assessment in a health facility. Our result indicated, by contrast, that only 71% of clinicians actually performed and documented a rabies risk assessment. The findings of this study suggest that risk assessment by clinicians and PEP decisions was mostly based on the type of exposure (i.e. bite, bite with puncture wound, licks, nibbles, indirect exposure) which are clearly outlined in the national guidelines (Table 1). However, they tend to misclassify the risk based on the patient’s answer. It is a concern that 13% of patients were mis-classified as having no risk, while they had moderate or severe risk.

Irrespective of the risk assessment, the vast majority of patients (91.6%) still received ARV treatment even when the risk was not assessed. Clinicians in high-risk areas of Bhutan thus proved relatively conservative in their attitude towards PEP prescription. However, for the 23 patients in the study who did not receive ARV, 16 were misclassified as having no rabies risk category, including three patients that were in fact in the severe risk category. Conversely, the only patient with no risk of rabies was still prescribed ARV, and another in the category of moderate risk was unduly prescribed RIG. This is similar to findings of a nationwide study conducted between 2005 and 2008, which reported frequent PEP administration in category I exposures [10]. The discrepancy between clinician’s practice and the national guidelines in our study lies in underestimating the rabies risk in the first place, rather than a lack of compliance with recommendations in subsequent PEP prescription. As a result, seven patients (2.6%) in the highest risk category had not received the appropriate treatment (neither ARV nor RIG). In this study, eight patients were exposed to “confirmed rabies” cases, all of which were cows (the exposure consisting of handling the carcass and drinking raw milk). Another 76 patients were exposed to animals classified as “suspected of rabies” by the clinician, including 62 dog bites. However, RIG was prescribed to only 3 patients (1%) which is in concurrence with the results of the earlier study that RIG was not regularly administered to dog bite victims in Bhutan [10]. Two of the patients receiving RIG were bitten by dogs suspected of rabies, the third after drinking milk and handling the carcass of a confirmed rabid cow. The risk for this patient was mis-classified by the clinician from moderate (according to the guidelines) to high. This highlights the need to raise the awareness of clinicians to more frequently prescribe RIG for patients bitten by suspected rabid animals, as the situation in this respect does not seem to have changed since 2011 [10]. In Bhutan, the availability of RIG is very limited, as in many rabies endemic countries [15]. Hence, the national guidelines contain modified recommendations indicating RIG for only most severe category 3 exposure, i.e. exposure to “suspected or confirmed” rabid cases. However, these terms are not clearly defined in the guidelines. For dog bites, discrepant advice co-exists to take into account extra-severity criteria, such as free-roaming versus pet dog, or vaccination status. Improving the supply in RIG treatment and clarifying the guidelines would help to improve the decisions on RIG by clinicians.

Clinical experience and good clinical judgment are essential to prevent human rabies [16]. Other studies on clinician’s knowledge and attitudes, conducted in the USA (Florida, Kentucky), showed an unsatisfactorily low level of compliance with national guidelines resulting in inappropriate PEP treatment [17-19]. Studies in countries free of rabies [20] or in endemic areas [21-23] similarly reported insufficient rabies risk assessment due to a lack of familiarity of clinicians with the recommendations, and highlighted the need for knowledge updating [14]. Lack of agreement between rabies risk assessment by clinicians and the guidelines was also apparent in our study, although only a low percentage resulted in subsequent inappropriate prescription. In contrast, public health physicians in Israel showed a very high level of compliance to PEP guidelines [24]. This could be due to different public health policies or better training or expertise of clinicians undertaking rabies assessment in Israel compared to clinicians in Bhutan. Moreover, access to immunoglobulin may not limit the compliance with the recommendations in Israel.

From a resource allocation standpoint, greater compliance with guidelines was shown to increase the benefit-cost ratio of PEP use in other studies [25, 26]. A study in a low rabies-risk area (Massachusetts, USA) highlighted how large amounts of rabies PEP could be wasted in patients with low or non-existent risk of rabies exposure [27]. By contrast, our study in high risk area showed that due to a tendency to underestimate the rabies risk, patients with moderate or severe risk were sometimes not prescribed PEP. A similar study in low rabies-risk areas of Bhutan would potentially shed light on potential differences in PEP decision patterns and whether unnecessary treatments occur more frequently in those areas.

This study highlighted important gaps in clinical practice in the management of patients exposed to rabies risk from animals in high-rabies-risk districts in Bhutan. Further progress in preventing human rabies deaths could be achieved by improving the clinical decisions in the event of animal exposure, while rationalizing the use of ARV and RIG. One key strategy is an ongoing education of clinicians to improve the accuracy of rabies risk assessment. An update of the national guidelines should accompany the training of professionals. In particular, improving clarity about epidemiological criteria for RIG prescription would help to prioritize patients needing RIG treatment. RIG administration in high-risk areas of the country, where category 3 exposure is frequent, is currently very low. In line with current WHO recommendation, optimizing access and uptake of RIG by animal bite victims through use of modified recommendations at the local level is strongly recommended including integrated management of animal bites patients in rabies endemic areas [4]. Further studies in low-rabies-risk areas of Bhutan are required for a full assessment of the situation regarding attitudes and practices of clinicians towards rabies in this country.

## Acknowledgments

We thank the following organization and the staffs whose contributions made this study possible. The Chief Medical Officer and focal health staff in 13 hospitals under Ministry of Health, Royal Government of Bhutan. The study team also is grateful to staff at the Faculty of Nursing and Public Health (FNPH), Khesar Gyalpo University of Medical Sciences of Bhutan and staff at the College of Natural Resources (CNR), Royal University of Bhutan.

## Supporting files

**S1 spreadsheet. National Rabies vaccine and immunoglobulin procurement record for final years 2014-15, 2015-16, 2016-17**

**S2 spreadsheet. dataset**

